# The neuroendocrine peptide catestatin promotes clearance of cutaneous *Staphylococcus aureus* through mast cell Mrgpr activation

**DOI:** 10.1101/2025.06.12.659211

**Authors:** Colin Guth, Hannah Dychtenberg, Erin Rudolph, Austin Pozniak, Sukhmeen Gill, Priyanka Pundir

## Abstract

Methicillin-resistant *Staphylococcus aureus* (MRSA) is a leading cause of cutaneous infections, underscoring the need for alternative therapeutic strategies. Catestatin, a neuroendocrine antimicrobial peptide produced by neurons and epithelial cells, has been implicated in skin defense against pathogens such as MRSA, though its mechanisms remain unclear. Here, we show that catestatin expression is upregulated in MRSA-infected skin wounds and that topical catestatin application significantly reduces MRSA burden in infected murine cutaneous wounds. This effect is dependent on the mast cell-specific G protein-coupled receptor Mrgprb2, the murine ortholog of human MRGPRX2. Notably, catestatin treatment leads to Mrgprb2-dependent suppression of inflammatory cytokine production and leukocyte infiltration, alongside upregulation of the antimicrobial peptide *Defb14*. In human mast cells, catestatin induces MRGPRX2-dependent degranulation, histamine release, prostaglandin D2 production, and cytokine expression. Pharmacological inhibition and western blot analysis reveal that catestatin activates multiple downstream G protein-dependent signaling pathways in an MRGPRX2-dependent manner. These findings demonstrate that catestatin promotes bacterial clearance by activating mast cells through Mrgprb2, thereby enhancing antimicrobial peptide production. Our study positions catestatin as a promising mast cell-targeting immunotherapeutic candidate for treating antibiotic-resistant skin infections.

## INTRODUCTION

Among all pathogens, *Staphylococcus aureus* is the leading cause of cutaneous infections, and the rise of methicillin-resistant *S. aureus* (MRSA) strains has amplified this threat^1–3^. Thus, alternative strategies to combat this pathogen are urgently needed. Host-derived antimicrobial peptides are a promising solution. In addition to direct antimicrobial activity, they also play immunomodulatory roles^4–6^, contributing to the elimination of skin pathogens^7–12^. Neuropeptides are another class of molecules recently shown to be critical in mediating bacterial infections^13–15^, suggesting another potential therapeutic avenue which utilizes endogenous mechanisms. One emerging peptide, catestatin, falls into both categories, defined as a neuroendocrine antimicrobial peptide, and has been shown to protect against cutaneous bacterial pathogens^16, 17^. However, the mechanisms underlying this effect remain unclear.

Catestatin is a cationic neuropeptide produced by both cholinergic and sensory neurons in the skin^18, 19^ and acts as a nicotine receptor antagonist, providing negative feedback to regulate cholinergic signalling^18^. Recently, catestatin has been discovered to influence physiological processes such as angiogenesis^20, 21^, hypertension^22–24^, cardiac function^24, 25^, metabolism^26^, and diabetes^27^, emphasizing a profound systemic effect which it often achieves via immunomodulation of macrophages, skewing them towards anti-inflammatory phenotypes^24, 27, 28^. In the context of the skin, catestatin is upregulated in both keratinocytes and neurons following skin barrier disruption and infection^16, 17^, improves symptoms of atopic dermatitis^29^ and promotes epidermal cell migration and proliferation in vitro^30, 31^. Additionally, despite relatively limited bactericidal activity in vitro^17^, mice lacking the catestatin precursor, chromogranin A, are more susceptible to MRSA and *Streptococcus pyogenes* infections^16^, suggesting an unexplored immunomodulatory role. Given the immunomodulatory effects of catestatin in other systems, it seems logical that it achieves its effects in the skin similarly; however, this is yet to be examined experimentally.

Mast cells are key candidates in this context, serving as tissue resident innate immune sentinels that rapidly respond to bacterial infections. Though best known for roles in allergy, mast cells also mediate pathogen defense and neuroimmune interactions^32–40^. Central to this is the G protein-coupled receptor MRGPRX2 (Mrgprb2 in mice)^41, 42^, expressed exclusively by connective tissue mast cells and activated by cationic ligands, including bacterial quorum-sensing molecules^33^, neuropeptides^41, 43, 44^, and antimicrobial peptides^5, 45^. Mast cells and neurons are closely associated in the skin, forming a functional unit^43, 46^, and Mrgprb2 can be activated by the neuropeptide substance P, mediating allergic responses^46^, neurogenic inflammation^43^, and pain^43^. Importantly, Mrgprb2-deficient mice show increased susceptibility to bacterial infections due to impaired quorum sensing interception^33^, and targeted Mrgprb2 activation improves infection outcomes in cutaneous models of infection^33, 39^.

While it is becoming increasingly clear that mast cells and specifically MRGPRX2/Mrgprb2 are essential for pathogen defense, the role of neuroimmune activation of mast cells through MRGPRX2/Mrgprb2 in this context remains unknown. Interestingly, catestatin has been shown to induce activation of mast cells in a G protein-dependent manner in vitro^47^, but the responsible receptor and the applicability of this relationship in vivo is unclear. Given the susceptibility of catestatin-deficient mice to MRSA infections despite low levels of bactericidal activity in vitro^16, 17^, and the influence over immune cells which catestatin exerts in other tissues^24, 27, 28^, we hypothesize that catestatin protects against cutaneous MRSA infections via immunomodulatory mechanisms. Further, given that catestatin can activate mast cells independently of the canonical catestatin receptor, and that it is a cationic (net charge at pH 7.0 = +4.9) neuroendocrine antimicrobial peptide^5, 41, 43, 45^, similar to other MRGPRX2/Mrgprb2 ligands, we hypothesize that catestatin exerts this immunomodulatory effect by activating mast cells via MRGPRX2/Mrgprb2. Here, we report that catestatin activates both mouse Mrgprb2 and human MRGPRX2. We demonstrate in vivo that application of catestatin to MRSA-infected cutaneous wounds enhances bacterial clearance via Mrgprb2. Interestingly, while we see an overall anti-inflammatory effect of catestatin, we also see an upregulation of the antimicrobial peptide β-defensin-14. In vitro, using a human mast cell line genetically modified to lack MRGPRX2, we show that catestatin induces robust MC activation via a variety of G protein-dependent signalling pathways. Taken together, these results reveal a neuroimmune mechanism which mediates the resolution of cutaneous wound infections.

## RESULTS

### Catestatin protects against bacterial infection in an Mrgprb2-dependent fashion

Since MRSA is the most commonly isolated pathogen from cutaneous wound infections^1^, we employed an infected full-thickness excisional wound model where a circular wound with a diameter of 6mm is created on the dorsum, penetrating fully through both the epidermis and dermis^48^ and then inoculated with 10^6^ CFUs of MRSA, strain USA300 LAC. It has been shown by others that catestatin and its precursor, chromogranin A, are upregulated following both infection and superficial tape-stripping injury, which damages the epidermis but leaves the dermis intact^17^. We sought to determine if the same held true in our wound infection model. While we saw no increase in gene expression of chromogranin A at 3 h post-injury in sterile wounds relative to uninjured skin (Fig. S1A), we did see chromogranin A gene expression upregulated 10-fold relative to uninjured skin in MRSA-infected wounds at 24 h post-infection (Fig. 1A), demonstrating that catestatin is upregulated following MRSA infection of cutaneous wounds. We next sought to determine whether topical application of exogenous catestatin improved bacterial clearance via immunomodulation. While catestatin is only directly bactericidal against MRSA at doses higher than 100 µM in vitro^17^, to mitigate any possible direct bactericidal effects we pre-treated wounds with either 50 µM of catestatin, or saline as a vehicle control, 1 h prior to infection. To assess the requirement of Mrgprb2 for any observed effects, we performed these experiments using both male and female wildtype and Mrgprb2-knockout (KO) C57Bl/6J mice, generated by McNeil et al^41^. Importantly, the Mrgprb2-KO mice are a functional knockout model, meaning that they are immunocompetent and still possess mast cells which are entirely functional other than their lack of Mrgprb2^41^. 24 h following inoculation, weight change, wound size and bacterial load were measured. In wildtype mice, catestatin-treated wounds showed a 45% reduction in bacterial load compared to vehicle-treated wildtype wounds (Fig. 1B), demonstrating that catestatin is effective in promoting MRSA clearance from wound infections. Additionally, we did not observe any difference in bacterial load between vehicle-treated and catestatin-treated wounds in Mrgprb2-KO mice (Fig. 1B), indicating that catestatin’s ability to enhance bacterial clearance is Mrgprb2-dependent. No changes were observed between any groups in wound size (Fig. 1C) or body weight change (Fig. S1B).

**Figure 1.**
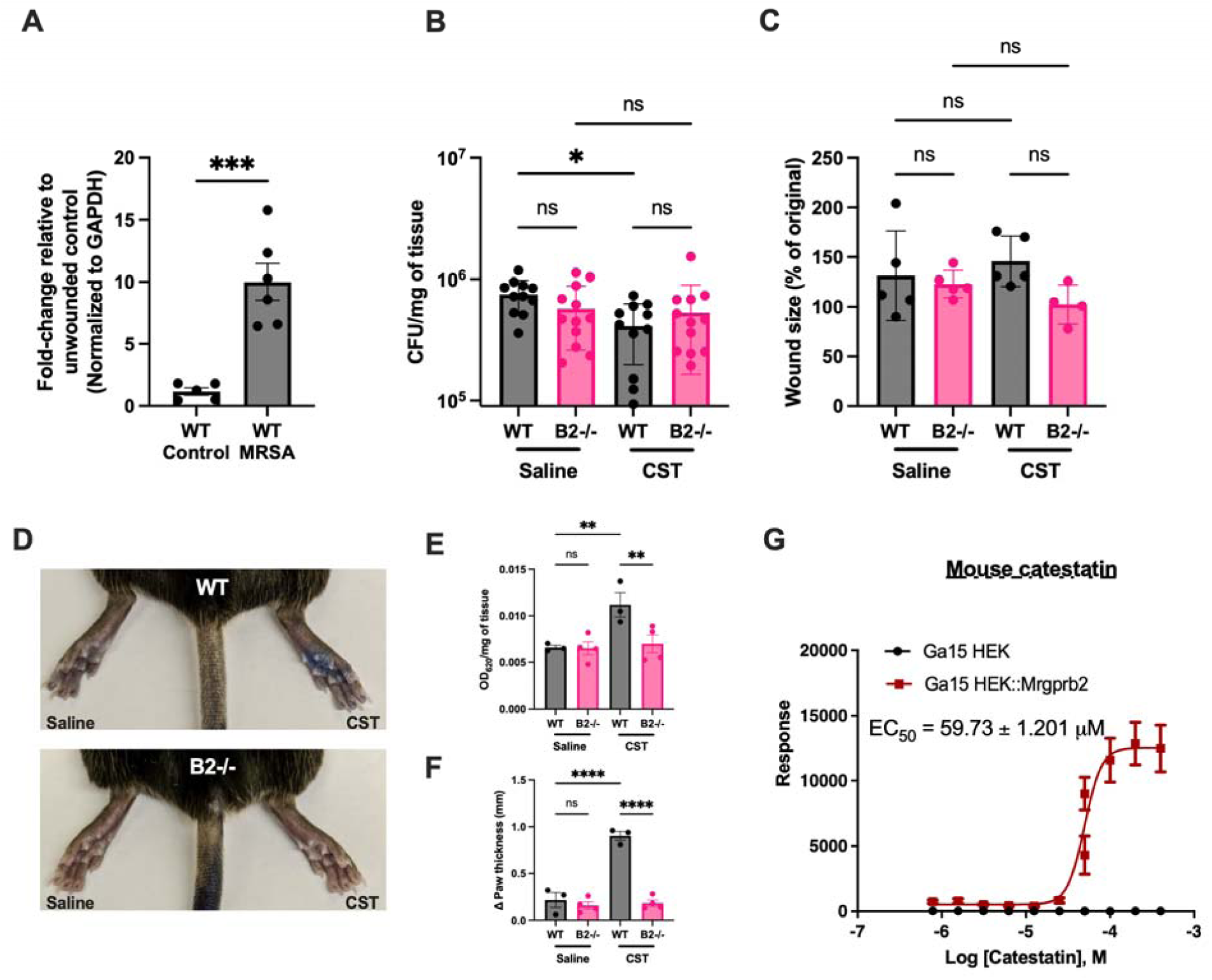
Treatment of wound infection with catestatin improves bacterial clearance in an Mrgprb2-dependent manner. **(A)** Gene expression levels of *CHGA* in healthy skin and skin from mice 24 h after MRSA infection, as determined by RT-qPCR. Data are normalized to *GAPDH* and expressed relative to mRNA levels in healthy skin samples. WT control, n=5; WT MRSA, n=6. **(B)** Quantification of colony-forming units (CFUs) recovered from MRSA-infected wounds of both wildtype (WT) and Mrgprb2-KO (B2-/-) C57BL/6J mice with and without catestatin (CST; 50 µM) treatment. Data are normalized to the weight of the wound sample. WT saline, n=11; WT CST, n=12; B2-/- saline, n=12; B2-/- CST, n=12. **(C)** Comparison of the change in wound size 24 h post-infection between saline-treated WT (n=5), CST-treated WT (n=5), saline-treated B2-/- (n=5) and CST-treated B2-/- mice (n=4). Data are expressed as the percentage of the area 24 h after infection relative to the area immediately after the wound was created. **(D)** Representative images corresponding to the extravasation assay in panel E. **(E and F)** WT and B2-/- mice were injected intravenously with Evans Blue. CST (50 µM) or saline were injected intradermally into the hindpaw. Dye extravasation quantified by OD_620_ normalized to the weight of each individual paw (E) and paw thickness measurements (F). WT, n=3; B2-/- n=4. **(G)** HEK cells expressing Mrgprb2 activation by CST (EC_50_=59.73 μM), as determined by FLIPR intracellular Ca^2+^ mobilization assay. n=4-9. Data are expressed as mean ± SEM, analyzed by unpaired Student’s *t*-test or one-way ANOVA. *P<0.05, **P<0.01, ***P<0.001, ****P<0.0001.

To further assess the specificity of catestatin for Mrgprb2 in the skin, we performed a passive cutaneous anaphylaxis assay where Evans blue dye is first injected intravenously followed by subcutaneous injection of either vehicle control or test compounds into the hind paw. Fifteen minutes post-paw injection, dye extravasation and changes in paw thickness are measured to determine the degree of local swelling^41^. From this experiment, we observed a significant increase in both dye leakage (Fig. 1D, E) and paw thickness (Fig. 1F) in catestatin-treated paws compared to vehicle-treated paws in wildtype mice. Conversely, the effect of catestatin on either dye leakage (Fig. 1D, E) or paw size (Fig. 1F) was completely abrogated in Mrgprb2-KO mice. Using a FLIPR intracellular Ca^2+^ mobilization assay^12, 33^, we found that mouse catestatin robustly activated Mrgprb2 with an EC_50_ of 59.73 μM, closely aligning with the dose used in our in vivo experiments (Fig. 1G). Overall, these results reveal that topical treatment of MRSA wound infections with catestatin is effective in aiding bacterial elimination, and this effect is dependent on the mast cell receptor Mrgprb2.

### Catestatin dampens inflammatory immune activation via Mrgprb2

We next aimed to investigate the molecular and cellular mechanisms underlying catestatin’s ability to reduce MRSA bacterial loads in wound infections via Mrgprb2. Given the role of mast cells in pro-inflammatory reactions in the skin^39, 43^, we hypothesized that catestatin induced the release of pro-inflammatory mediators and promoted recruitment of leukocytes to the site of infection by activating Mrgprb2 on mast cells. To investigate this, we performed ELISAs targeting C-C motif ligand 2 (CCL2), tumor necrosis factor-α (TNF-α) and interleukin (IL)-6, common mast cell-derived mediators of cutaneous inflammation which promote early recruitment of leukocytes, including monocytes and neutrophils^37, 43^. Interestingly, contrary to our initial hypothesis, we observed a reduction in CCL2 (97.71 pg/mg of skin; Fig. 2A) and TNF-α (3.15 pg/mg of skin; Fig. 2B) in catestatin-treated wildtype wound infections, relative to vehicle-treated wildtype wound infections (CCL2=137.0 pg/mg of skin, TNF-α=5.46 pg/mg of skin; Figure 2A, B), and a trend towards reduction for IL-6, with 279.0 pg/mg of skin in catestatin-treated wildtype wounds and 385.60 pg/mg of skin in vehicle-treated wildtype wounds (Fig. 2C). No differences in cytokine concentrations were observed between vehicle- and catestatin-treated wound infections from Mrgprb2-KO mice (Fig. 2A-C), suggesting this effect is Mrgprb2-dependent.

**Figure 2.**
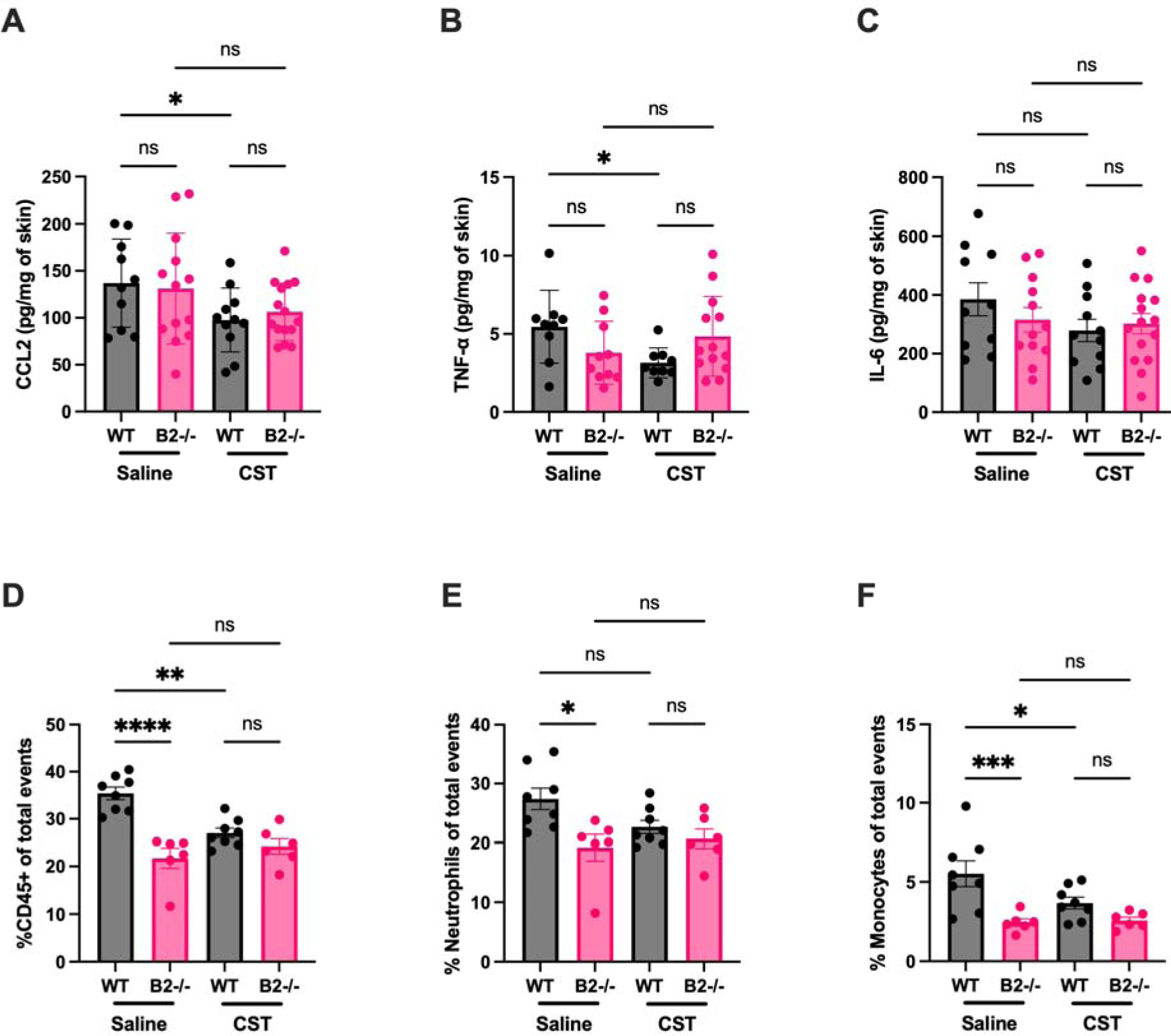
Catestatin dampens the inflammatory immune response to MRSA infection via Mrgprb2. **(A-C)** ELISA quantification of CCL2 (A), TNF-α (B), and IL-6 (C) levels in the skin from catestatin (CST; 50 μM) and saline-treated MRSA wound infections in wildtype (WT) and Mrgprb2-KO (B2-/-) mice 24 h post-infection. Data are normalized to the weight of the wound sample. WT saline, n=10; WT CST, n=11; B2-/- saline, n=13; B2-/- CST, n=15. **(D-F)** Flow cytometric quantification of immune cells in MRSA-infected skin from WT and B2-/- mice 24 h post-infection following treatment with CST (50 μM) and saline. Total CD45^+^ cells (D), neutrophils (Ly6C^+^Ly6G^+^; E), and monocytes (Ly6C^+^Ly6G^-^; F) are expressed as a percentage of total events. WT saline, n=8; WT CST, n=8; B2-/- saline, n=5; B2-/- CST, n=5. Data are expressed as mean ± SEM, analyzed by one-way ANOVA. *P<0.05, **P<0.01, ***P<0.001, ****P<0.0001.

ELISA results prompted us to investigate the effect of catestatin on recruitment of leukocytes, particularly neutrophils and monocytes which are both key mediators of the acute response to both tissue injury and infection^49–51^. Consistent with our ELISA results, relative to vehicle-treated controls, catestatin-treated wildtype mice showed a significant reduction in CD45^+^ cells, comprising 27.00% of live cells in catestatin-treated wildtype wound infections compared to 35.86% in vehicle-treated wildtype wound infections (Fig. 2D; Fig. S2). Monocytes, defined as Ly6C^+^Ly6G^-^, gated on live CD45^+^CD11b^+^ cells (Fig. S2), were also reduced in catestatin-treated wildtype wounds, with 3.67% compared to 5.52% in vehicle-treated wildtype wounds (Fig. 2F). We also observed a trend towards reduction of neutrophils, defined as Ly6C^+^Ly6G^+^, gated on live CD45^+^CD11b^+^ cells (Fig. S2), at 22.64% in catestatin-treated wildtype compared to 27.40% in vehicle-treated wildtype (Fig. 2E). No difference in any immune cell population was observed between vehicle- and catestatin-treated wound infections in Mrgprb2-KO mice (Fig.2 D-F). Healthy skin from wildtype and Mrgprb2-KO mice also showed no differences in cytokines or immune cell populations (data not shown).

### Catestatin induces expression of antimicrobial peptides via Mrgprb2 in MRSA-infected wounds

Both catestatin and mast cells have been shown to drive an increase in the production of a variety of other antimicrobial peptides in the skin, namely cathelicidin, β-defensins and S100A7 (also known as psoriasin)^16, 37^. Recent studies using robust wound infection models show that mast cells promote bacterial clearance not by recruiting leukocytes, but by stimulating antimicrobial peptide release from keratinocytes^37^. Thus, we sought to determine if catestatin promotes antimicrobial peptide production via Mrgprb2, which could subsequently lead to a reduction in bacterial load in our MRSA wound infection model. Indeed, via RT-qPCR, we observed a 2.5-fold increase in *Defb14* gene expression in catestatin-treated wildtype wound infections relative to vehicle-treated wildtype wound infections, along with no change in expression between catestatin- and vehicle-treated Mrgprb2-KO wounds (Fig. 3A), once again suggesting that this effect is Mrgprb2-dependent. No change was observed for *Camp*, the gene responsible for cathelicidin production, or *S100a7,* the gene responsible for psoriasin, between any of the four groups (Fig. 3B, C).

**Figure 3.**
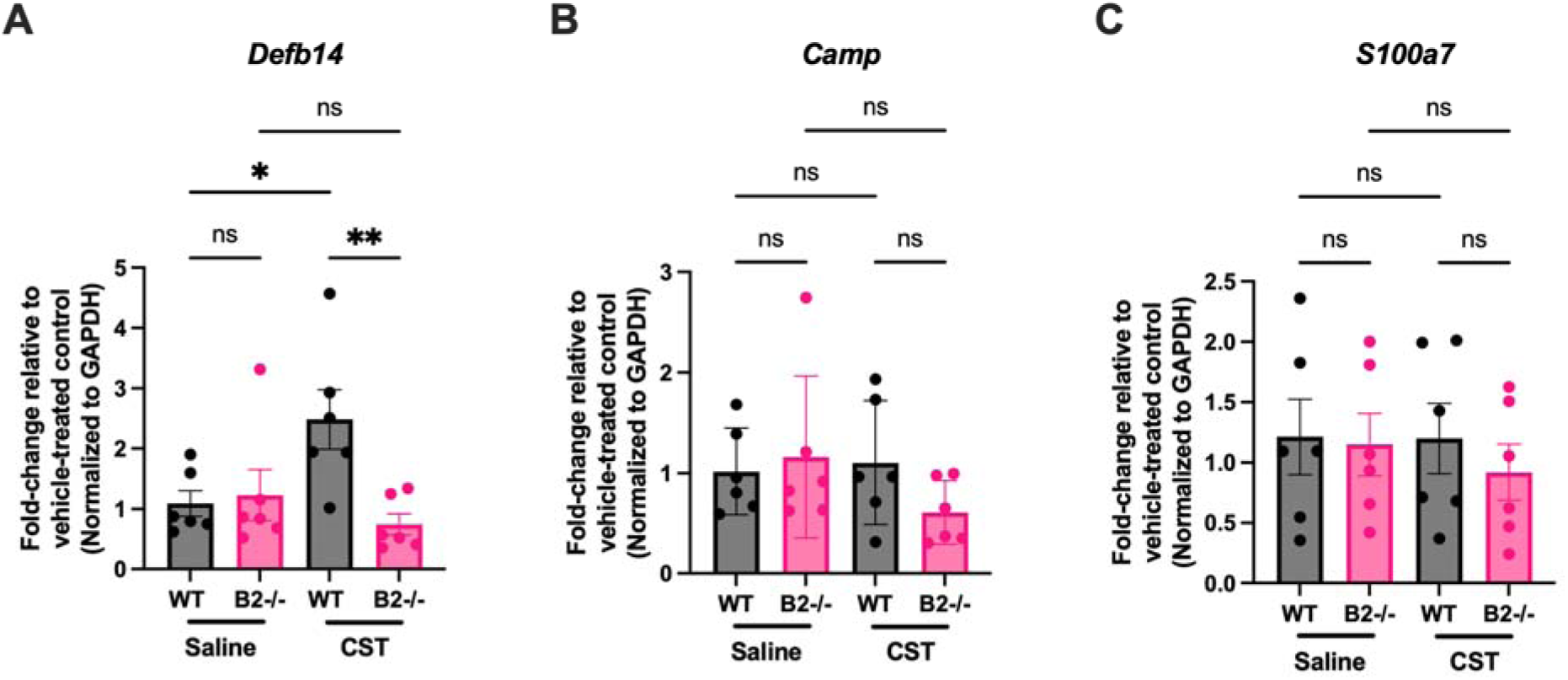
Catestatin induces expression of the antimicrobial peptide *Defb14* in MRSA-infected wounds via Mrgprb2. **(A-C)** Gene expression levels of *Defb14* (A), *Camp* (B), and *S100a7* (C) in skin from wildtype (WT) and Mrgprb2-KO (B2-/-) mice 24 h after MRSA infection, as determined by RT-qPCR. Data are normalized to *GAPDH* and expressed relative to mRNA levels in vehicle-treated control samples of the corresponding genotype. WT saline, n=6; WT CST, n=6; B2-/- saline, n=6; B2-/- CST, n=6. Data are expressed as mean ± SEM, analyzed by one-way ANOVA. *P<0.05, **P<0.01.

### Catestatin induces human mast cell degranulation in an MRGPRX2-dependent manner

Our in vivo results clearly demonstrated that the effect of catestatin depends on the murine mast cell receptor Mrgprb2, and that we can leverage this interaction to clear MRSA from wound infections. While it has been shown that catestatin activates human mast cells independently of the canonical catestatin receptor^47^, the receptor which mediates this effect is unknown. Thus, we aimed to determine if catestatin served as a ligand for Mrgprb2 and its human homologue, MRGPRX2. To test this hypothesis, we utilized the LAD2 human mast cell line, which expresses MRGPRX2^5^, along with MRGPRX2-knockout (X2-KO) LAD2 cells in which the expression of MRGPRX2 was silenced using CRISPR^52^. Rapid degranulation is core to the sentinel function of mast cells^32^. Thus, we began by measuring the release of the common marker of human mast cell degranulation, β-hexosaminidase^53^, from both wildtype and X2-KO LAD2 cells following treatment with catestatin. Additionally, as a positive control, cells were treated with a well-known MRGPRX2 agonist, compound 48/80^41, 54^. We observed dose-dependent release of β-hexosaminidase from wildtype but not X2-KO LAD2 cells in response to catestatin, with maximum release (net release 37.85%) at 10 µM, comparable to c48/80 (net release 44.58%; Fig. 4A). Functionally, the primary mediator within mast cell granules which produces the vascular leakage associated with mast cell degranulation is histamine^55^. Thus, we performed an assay to confirm that catestatin-induced degranulation also resulted in histamine release. Indeed, catestatin induced a dose-dependent release of histamine from wildtype LAD2s, with maximum release (408 ng/ml) at 10 µM, and this was again comparable to c48/80 (397 ng/ml), while this effect was absent in X2-KO LAD2s (Fig. 4B). To determine whether the lack of activation of our X2-KO cells was due to a refractory effect, we stimulated both wildtype and X2-KO cells through the canonical pathway of mast cell activation mediated by the FcεRI receptor^56^, and verified β-hexosaminidase release by both wildtype and X2-KO cells (Fig. S3A). Furthermore, we determined catestatin to not be cytotoxic by measurement of lactate dehydrogenase (LDH) release following catestatin treatment, where we saw no increase in LDH compared to negative control (Fig. S3B).

**Figure 4.**
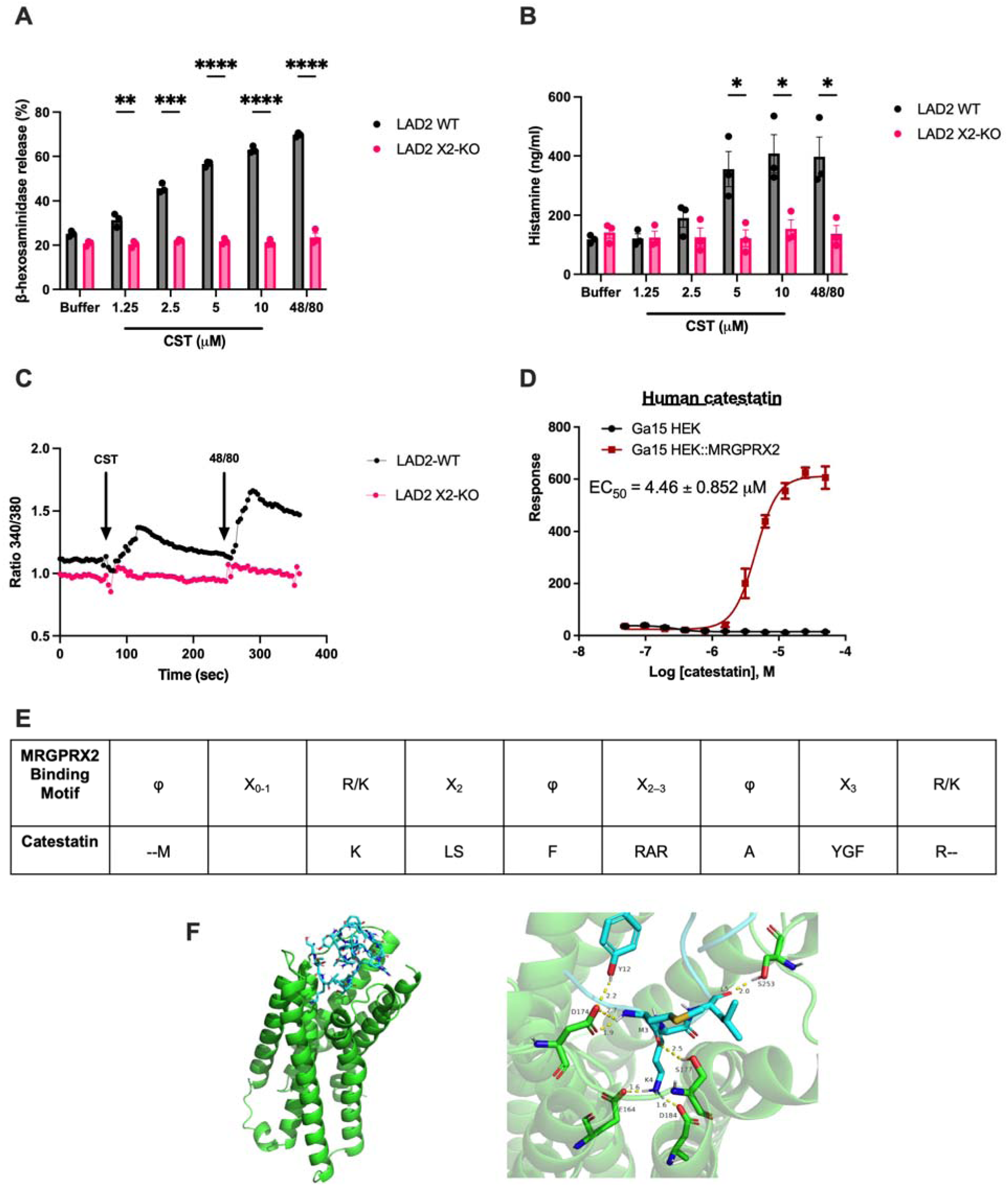
Catestatin induces human mast cell degranulation and calcium mobilization via MRGPRX2. **(A)** β-hexosaminidase release from wildtype (LAD2 WT) and MRGPRX2-KO (LAD2 X2-KO) LAD2 cells in response to increasing concentrations of catestatin and 0.5 µg/ml 48/80. n=3. **(B)** Histamine release from WT and X2-KO LAD2 cells in response to increasing concentrations of catestatin and 0.5 µg/ml 48/80. n=3. **(C)** Intracellular Ca^2+^ mobilization in Fura-2-loaded WT and X2-KO LAD2 cells following stimulation with 10 μM catestatin (CST) at 70 s and 0.5 μg/ml 48/80 at 256 s. n=3. **(D)** HEK cells expressing MRGPRX2 activation by CST (EC_50_=4.46 μM), as determined by FLIPR intracellular Ca^2+^ mobilization assay. n=4-9. **(E)** Alignment of a portion of the catestatin peptide sequence with the MRGPRX2 ligand binding motif identified by Yang et al^47^. **(F)** Visualizations generated using PyMOL of molecular docking simulations between MRGPRX2 (green) and catestatin (blue) performed using HADDOCK. Residues are labelled using single-letter amino acid codes and hydrogen bonds (yellow) are shown between interacting residues, labelled with the length of the bond in angstroms. Data are expressed as mean ± SEM, analyzed by unpaired Student’s *t*-test. *P<0.05, **P<0.01, ***P<0.001, ****P<0.0001.

Calcium mobilization is essential for GPCR-driven mast cell degranulation^57^. To investigate the mechanism underlying catestatin-induced degranulation in LAD2 cells, we assessed Ca^2+^ flux using a Fura-2 assay. Catestatin (10 μM) elicited a Ca^2+^ response, with compound 48/80 as a positive control (Fig. 4C). Notably, the effect was abolished entirely in X2-KO cells (Fig. 4C). Furthermore, using HEK293 cells expressing MRGPRX2, we found that catestatin activated the receptor with an EC50 of 4.47 μM (Fig. 4D). Altogether, these results demonstrate that catestatin induces degranulation and histamine release from human mast cells in an MRGPRX2-dependent manner.

Recently, the structural elements involved in MRGPRX2-ligand binding have been determined via cryo-electron microscopy (cryo-EM) studies, revealing that MRGPRX2 binds peptide ligands via a hydrophobic region near the opening of the binding site and an acidic region located closer to the core of the receptor^58^. This study also revealed a peptide binding motif (φ^p9^(X_0–1_) R/K^p10^(X_2_) φ^p13^(X_2–3_) φ^p16^(X_3_) R/K^p20^) present in a variety of known MRGPRX2 ligands^58^, which catestatin possesses as well (Fig. 4E). Utilizing the structure of MRGPRX2 bound to the peptide ligand cortsitatin-14^58^, as well as a nuclear magnetic resonance structure of catestatin^59^, we performed molecular docking analysis to predict the binding of catestatin with MRGPRX2 using High Ambiguity Driven protein-protein DOCKing (HADDOCK)^60, 61^. As reference interacting residues for catestatin, we used both the hydrophobic and basic residues within the binding motif (Met3, Lys4, Leu5, Phe7, Arg8, Ala9, Arg10, Ala11, Tyr12, Phe14, Arg15) and for MRGPRX2 we used the acidic and hydrophobic residues shown to comprise the binding site of MRGPRX2 (Glu164, Phe170, Asp184, Phe239, Trp243, Asp252, Asp254, Phe257, His259)^58^. Similar to interactions between MRGPRX2 and other ligands, this modelling predicted hydrogen bonding interactions shorter than 3.0 Å in length between basic and hydrophobic residues of catestatin with acidic and hydrophobic residues of MRGPRX2 (Fig. 4F). Particularly, interactions between catestatin Lys4 and the primary acidic residues comprising the MRGPRX2 binding pocket, Glu164 and Asp184, were observed, both forming hydrogen bonds with lengths of 1.6 Å (Fig. 4F). Similar to other peptide ligands of MRGPRX2, we also observed a variety of hydrophobic interactions between MRGPRX2 and catestatin are shown as well (Fig. 4F). This modelling provides further evidence of catestatin as an MRGPRX2 ligand, demonstrating similarities in binding with other known ligands.

### Catestatin induces early-phase lipid mediator and late-phase cytokine release in an MRGPRX2-dependent manner

While degranulation is an essential property of mast cell activation, they also release a variety of other mediators. Namely, lipid mediators such as prostaglandins and cysteinyl leukotrienes, as well as a variety of cytokines, chemokines and growth factors^62, 63^. Thus, we aimed to determine if catestatin-induced mast cell activation via MRGPRX2 only led to mast cell degranulation or a more robust response. First, we measured release of the lipid mediator prostaglandin D_2_ (PGD_2_), as mast cell-derived PGD_2_ is known to play important roles in orchestrating immune responses in the skin^64^. Similar to degranulation, we found that catestatin at 10 µM induces PGD2 release (292.11 pg/ml; Fig. 5A), revealing that catestatin induces lipid mediator release from mast cells in an MRGPRX2-dependent fashion. Next, we performed RT-qPCR to compare gene expression of the cytokines TNF-α, CCL2, IL-6, IL-3 and granulocyte-macrophage colony-stimulating factor (GM-CSF) between wildtype and X2-KO cells following catestatin treatment. In catestatin-treated wildtype cells, we observed a greater than two-fold increase in cytokine gene expression compared to untreated controls for all targets, whereas X2-KO cells showed no change (Fig. 5B-F). We then followed up on these changes in gene expression by measuring protein release of CCL2 and TNF-α via ELISA and observed a significant increase in CCL2 (145 pg/ml) following treatment with 10 µM catestatin in wildtype but not X2-KO cells (75 pg/ml; Fig. 5G). However, despite an increase in gene expression, TNF-α release was not induced by catestatin, even as high as 20 µM (Fig. 5H). The production of GM-CSF was also not induced upon catestatin treatment of LAD2 cells (Fig. 5I). Altogether, these results display that catestatin induces robust activation and release of early lipid mediators and cytokines from human mast cells in an MRGPRX2-dependent fashion.

**Figure 5.**
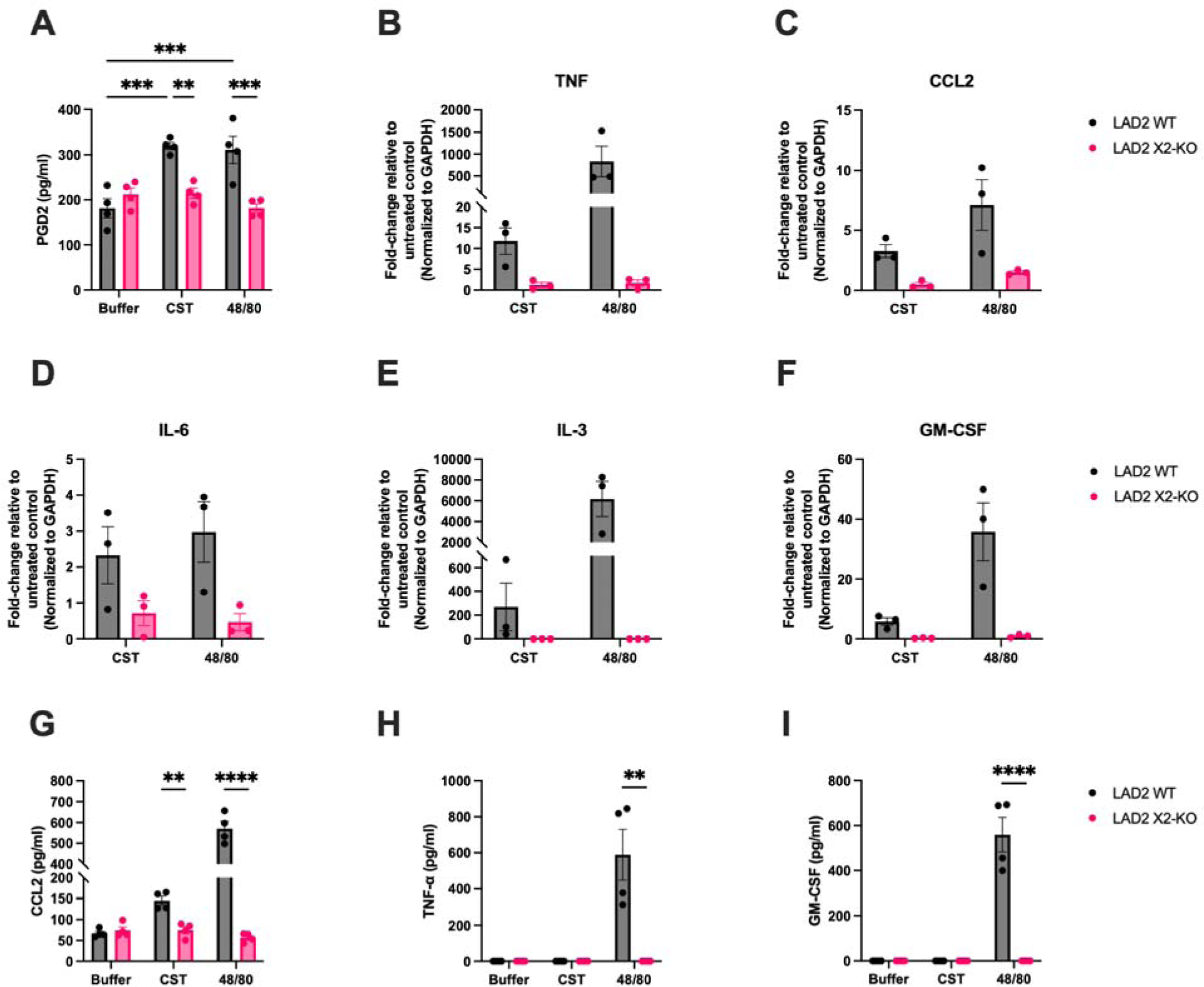
Catestatin induces *de novo* mediator release from human mast cells via MRGPRX2. **(A)** Prostaglandin D_2_ (PGD_2_) release from wildtype (LAD2 WT) and MRGPRX2-KO (LAD2 X2-KO) LAD2 cells in response to 10 µM catestatin (CST) and 0.5 µg/ml compound 48/80. n=4 **(B-F)** Gene expression of TNF-α (B), CCL2 (C), IL-6 (D), IL-3 (E), and GM-CSF (F) in WT and X2-KO LAD2 cells in response to CST (10 µM) and 48/80 (0.5 µg/ml). All data are normalized to GAPDH and expressed relative to untreated control samples. n=4. (**G-I**) Protein production of CCL2 (G), TNF-α (H), and GM-CSF (I) release from WT and X2-KO LAD2 cells stimulated with CST (10 µM for CCL2 and GM-CSF, 20 µM for TNF-α) and 48/80 (0.5 µg/ml for CCL2 and GM-CSF, 1 µg/ml for TNF-α). n=4. Data are expressed as mean ± SEM, analyzed by unpaired Student’s *t*-test. *P<0.05, **P<0.01, ***P<0.001, ****P<0.0001.

### Catestatin activates a robust signalling cascade downstream of MRGPRX2

To further investigate the underlying mechanisms of catestatin-induced mast cell activation, we interrogated the activation of signalling proteins downstream of MRGPRX2 following catestatin treatment. Chemical inhibition of phospholipase C (PLC; Ro-31-8220), protein kinase C (PKC; U-73122) and phosphatidylinositol 3-kinase (PI3K; wortmannin), signalling pathways revealed that catestatin-induced mast cell degranulation was sensitive to all inhibitors tested (Fig. 6A-C), indicating catestatin induces a robust signalling cascade involving many G protein-dependent pathways to achieve degranulation. We then analyzed activation of the MAPK protein ERK1/2, which is downstream of the previously examined signalling proteins and is known to play a role in mast cell activation^65^ in wildtype and X2-KO cells via western blot by measuring both unphosphorylated, or native, ERK1/2 in addition to phosphorylated ERK1/2 to determine the degree of activation. A time course analysis was first performed to determine the optimal timepoint for catestatin-induced ERK1/2 activation, which revealed peak phosphorylation at 5 min post-treatment (Fig. S4). Using this optimized time point, we then treated both wildtype and X2-KO cells with catestatin and examined ERK1/2 phosphorylation. While catestatin increased ERK1/2 phosphorylation in wildtype cells, no phosphorylation was observed in X2-KO cells (Fig. 6D). Thus, we conclude that catestatin stimulates mast cell activation through distinct G protein signalling pathways downstream of MRGPRX2.

**Figure 6.**
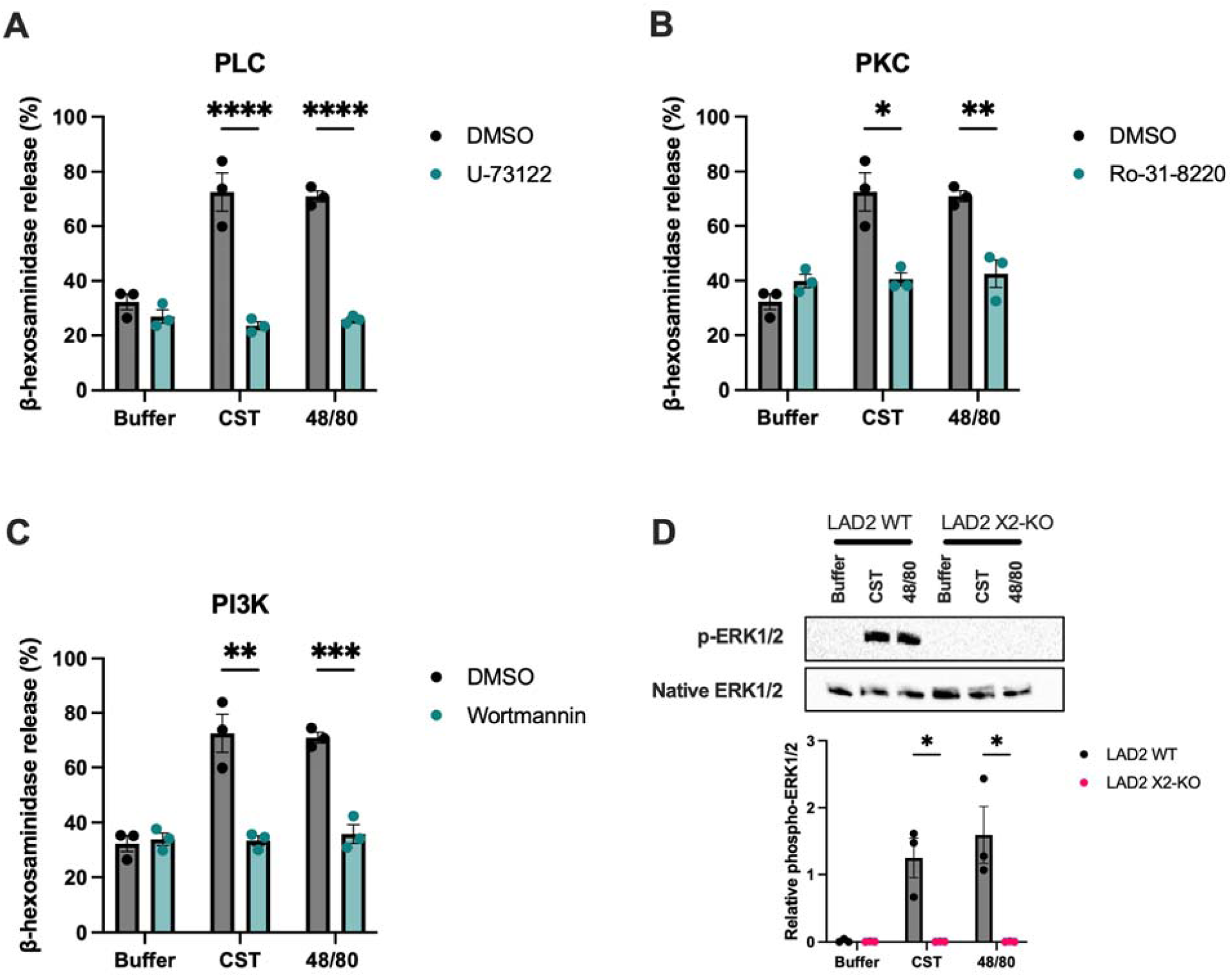
Catestatin activation of mast cells is dependent upon multiple signaling pathways. **(A-C)** Effect of pharmacological inhibition on catestatin (CST; 10 µM) and 48/80 (0.5 µg/ml)-induced degranulation of wildtype (LAD2 WT) and MRGPRX2-KO (LAD2 X2-KO) LAD2 cells. Cells were pre-treated with U-73122 (PLC inhibitor; 10 µM; A), Ro-31-8220 (PKC inhibitor; 10 µM; B), and Wortmannin (PI3K inhibitor; 10 µM; C) prior to stimulation. n=3. **(D)** Western blot analysis of ERK1/2 phosphorylation in LAD2 WT and LAD2 X2-KO cells in response to catestatin (10 µM) or compound 48/80 (0.5 µg/ml). Amount of ERK1/2 phosphorylation was normalized to the levels of native ERK1/2 present in the sample. n=3. Data are expressed as mean ± SEM, analyzed by unpaired Student’s *t*-test. *P<0.05, **P<0.01, ***P<0.001, ****P<0.0001.

## DISCUSSION

Neuroimmune mechanisms regulate a variety of homeostatic processes such as social behaviours65^66–69^, sleep^70–72^, and interactions along the gut-brain axis^73, 74^. More recently, a role for neuroimmune interactions between peripheral sensory neurons and resident immune cells has been shown to mediate bacterial infections^13–15^. Understanding the bidirectional communication between the immune and nervous systems, especially as they relate to disease, presents opportunity not only to gain insight into the mechanisms by which the body defends itself, but also to leverage these mechanisms to design therapeutics which bolster endogenous defense systems. One limitation of the field as it currently stands in relation to bacterial infections is a focus on only a few neuropeptides, namely calcitonin gene-related peptide and substance P. Thus, we aimed to explore the role of a relatively understudied neuropeptide, catestatin, as it relates to neuroimmune defense against bacterial pathogens.

Initially discovered as a component of the neuroendocrine system, catestatin is released by cholinergic and sensory neurons and acts as a nicotinic receptor antagonist^18, 19^. Recently, catestatin has been discovered to provide a protective role in the context of cardiac dysfunction^24^, gastrointestinal inflammation^28^ and diabetes^27^. In these contexts, catestatin promotes macrophage polarization towards an anti-inflammatory phenotype, mitigating damage caused by excessive inflammation^24, 27, 28^. Interestingly, catestatin has been correlated with atopic dermatitis in humans^29^. Affected individuals exhibit lower levels of its precursor, chromogranin A, and catestatin improves dermatitis symptoms in mouse models by suppressing Th2 cytokines (IL-4, IL-5, and IL-13) and promoting skin integrity proteins from keratinocytes^29^. Catestatin has also been shown to mediate bacterial infections in the skin^16, 17^. Catestatin is upregulated in the skin following disruption to the skin barrier as well as bacterial infection, and mice lacking the catestatin precursor chromogranin A are more susceptible to MRSA and *Streptococcus pyogenes* infections^16^. While two of these studies in the skin have shown that catestatin could exert effects on keratinocytes^16, 29^, the roles of immune cells in catestatin-mediated bacterial defense were not explored. Here, we reveal that catestatin indeed influences innate immunity to control bacterial skin infections.

Using a MRSA-infected cutaneous wound model in mice, we found that catestatin treatment significantly reduces bacterial burden 24 h post-infection. We hypothesized that this effect is mediated through the mast cell-expressed receptor Mrgprb2, which responds to ligands with similar properties to catestatin. Indeed, in Mrgprb2-deficient mice, catestatin had no effect, revealing a critical immunomodulatory mechanism for catestatin’s function in bacterial defense. Somewhat unsurprisingly given the previously discussed reports in the literature of catestatin’s anti-inflammatory effects in other tissues, including the skin^24, 27–29^, we found that catestatin treatment induced immunosuppression, evidenced by a reduction in pro-inflammatory cytokines as well as leukocyte infiltration, and these effects were also Mrgprb2-dependent. It should be noted that significantly fewer leukocytes were observed in vehicle-treated Mrgprb2-KO wound infections compared to vehicle-treated wildtype wound infections. Despite this, we still observed no difference in the proportions of CD45^+^ cells, neutrophils, or monocytes between vehicle-treated Mrgprb2-KO wound infections and catestatin-treated Mrgprb2-KO wound infections, reinforcing the dependency of catestatin on Mrgprb2. Furthermore, we have observed no differences in immune profiles in the healthy skin between wildtype and Mrgprb2-KO mice, as also reported by others^43, 46^.

This immunosuppressive effect, though consistent with previous literature, raised the question: how does bacterial clearance still occur? Especially considering that immunosuppression tends to favour bacterial growth, with certain microbes purposefully inducing immunosuppression via neuroimmune circuits to facilitate invasion^14, 15^, these two results appeared contradictory. However, catestatin has been shown to prevent nicotine-induced suppression of antimicrobial peptides in the skin during infection^16^. Furthermore, in a model of *Pseudomonas aeruginosa* wound infection, mast cells were essential for bacterial clearance via induction of antimicrobial peptides from keratinocytes, independent of leukocyte recruitment^37^. Similarly, in our MRSA infected wound model, catestatin increased expression of the antimicrobial peptide *Defb14* in an Mrgprb2-dependent manner. Importantly, *Defb14* promotes clearance of MRSA in other skin infection models^12^, and lower levels of its human ortholog are associated with worse MRSA outcomes in the clinic^75^. While future experiments, such as *Defb14* knockout, or utilization of a blocking antibody, are needed to directly confirm Defb14’s role in catestatin-mediated clearance, our data support this as a likely mechanism. Together with the work of Zimmerman et al.^37^, these findings suggest that, in wound infections, mast cells promote bacterial clearance by inducing antimicrobial peptides rather than inflammation. This would provide a logical benefit to the host since mitigating inflammation is essential to proper wound healing^49, 51, 76^.

We also found that catestatin-induced immunosuppression is Mrgprb2-dependent, despite mast cells traditionally being considered pro-inflammatory^33, 39^. However, evidence increasingly shows mast cells can also resolve inflammation. For example, they are involved in resolving inflammation in response to biomaterial implantation via macrophage polarization and suppression of inflammatory cytokines^77^, in addition to immunosuppressive effects of mast cells induced by sympathetic neurons in the context of wound healing under psychological stress^78^, among other examples^79–84^. From a therapeutic perspective, a peptide that both clears bacteria and limits inflammation would be advantageous, as dampening inflammation promotes wound healing^51^. Our findings reveal a neuroimmune circuit in which catestatin activates mast cells via Mrgprb2, suppressing inflammation while enhancing antimicrobial peptide production, leading to reduced bacterial burden (Fig. 7).

**Figure 7.**
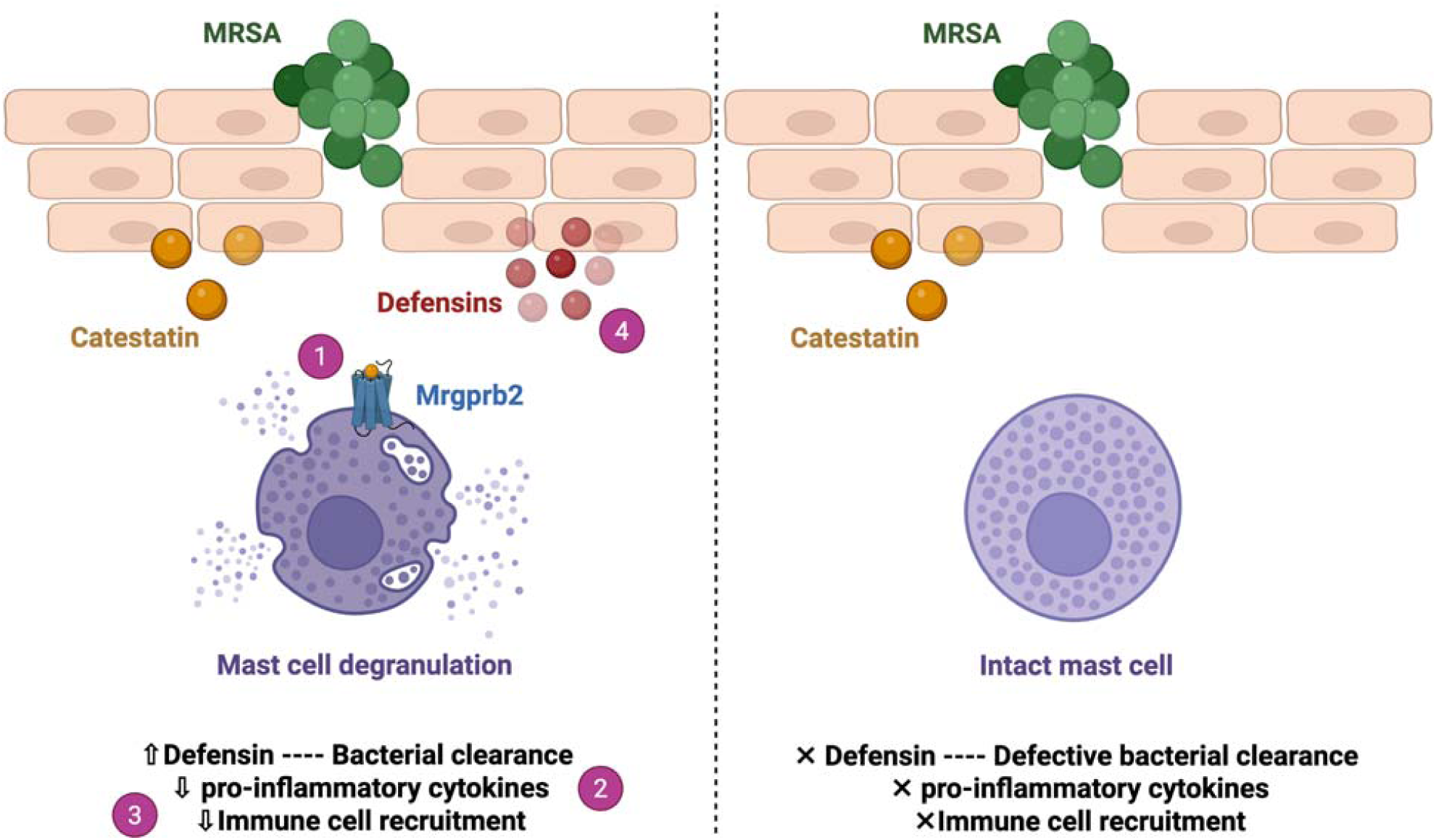
Graphical depiction of the proposed mechanism by which catestatin reduces bacterial burden in MRSA wound infections via mast cells and Mrgprb2. Catestatin activates mast cells via Mrgprb2 (1). While this activation leads to a net anti-inflammatory effect in the skin, characterized by lower cytokine levels (2) and diminished leukocyte recruitment (3), it also induces the upregulation of antimicrobial peptides (4), likely from epithelial cells, which have known bactericidal efficacy against MRSA. Created in https://BioRender.com.

Catestatin is shown to be present in skin and produced by keratinocytes and sensory neurons^17, 19^, with aberrant expression during injury, infection, and inflammation^17, 19, 29^. In our model, we observed upregulation of chromogranin A at 24 h post-MRSA infection but not at 3 h post-injury in sterile wounds. Radek et al. previously showed that tape-stripping, which superficially disrupts the epidermis but not the dermis, induced catestatin production, but was blocked by occlusion^17^. This suggests catestatin production may require extensive epidermal damage, exposing a large area of the stratum granulosum to air. In contrast, excisional wounds may not sufficiently expose this layer. Thus, exposure of the stratum granulosum may be necessary for catestatin induction at early timepoints following injury in the absence of a pathogen.

In 2011, Aung et al. demonstrated that human mast cells are activated by the neuroendocrine antimicrobial peptide catestatin^47^. Further, they revealed that human mast cells express the canonical catestatin receptor, but silencing of this receptor did not prevent catestatin-induced mast cell activation, leaving the mast cell receptor unknown^47^. Given catestatin’s structural similarity to known MRGPRX2 ligands^5, 41, 43, 44, 46, 85^, we hypothesized that MRGPRX2 is the human receptor. Here, we saw similar results to Aung et al., that catestatin induces robust activation of mast cells, eliciting the release of all three classes of mast cell mediators. Importantly, our experiments also utilized MRGPRX2-KO mast cells, for which the effects of catestatin were abolished, conclusively demonstrating that MRGPRX2 is the human mast cell receptor for catestatin. Further exploring the effect of catestatin on human mast cells, we also saw sensitivity of catestatin-induced mast cell activation to inhibition of a variety of signalling protein inhibitors, and that following catestatin treatment, the MAPK signalling pathway became activated.

In contrast to our in vivo results which showed an overall reduction in pro-inflammatory cytokines in catestatin-treated WT MRSA wound infections, we saw production of a variety of pro-inflammatory mediators induced by catestatin in vitro. Since the ELISAs we performed on wound samples look at the total amount of each cytokine produced by all cells in the wound sample, it is possible that mast cells do in fact upregulate these mediators in response to catestatin in vivo, but that the expression of these mediators by other cell types are driven down, resulting in a net reduction in each mediator in vivo. It is also possible that other mediators present in our wound infection model, which are not present in vitro, alter the response of mast cells to catestatin.

The rise of antimicrobial-resistant pathogens, particularly MRSA, poses a serious health threat^1, 86^. As traditional antibiotics lose efficacy, immune-based therapies offer promising alternatives. Enhancing endogenous immunity is a key principle in cancer immunotherapy and is now being applied to infectious diseases. Here, we show that exogenous application of the neuroendocrine antimicrobial peptide catestatin reduces MRSA burden in a murine infected wound model via Mrgprb2-mediated mast cell activation, resulting in reduced inflammation but increased antimicrobial peptide production. Furthermore, we establish MRGPRX2, the human homologue of Mrgprb2^41^, as the human mast cell receptor for catestatin. Our findings highlight a novel neuroimmune circuit that could be therapeutically harnessed to treat antibiotic-resistant infections.

## METHODS

### Animals

All procedures were performed in accordance with the guidelines set by the Canadian Council for Animal Care and all protocols (AUP 4806 and 4884) were approved by the University of Guelph Animal Care Committee. Male and female wildtype (WT) and Mrgprb2-KO (B2-/) C57BL/6J mice were used at 8-10 weeks of age for wound healing experiments and up to 6 months for all other experiments. Mice were housed in individually ventilated cages, with two to four mice per cage, in a Containment Level 2 (CL2) facility under a 12-hour light/dark cycle, with ad libitum access to food and water. Routine health surveillance confirmed that animals were free of all screened pathogens. Mice were monitored daily by animal care and laboratory staff, and no unexpected adverse events or mortality were observed during the study.

### Bacterial strain and growth conditions

*Staphylococcus aureus* CA-MRSA strain USA300 Lac was kindly provided by Dr. Georgina Cox (University of Guelph). To prepare inoculum for infectious models, cultures were streaked from glycerol stocks onto tryptic soy agar plates and incubated overnight at 37°C in a humidified chamber with 5% CO_2_. The next day, the culture was resuspended in tryptic soy broth (TSB) and added to 5 mL fresh TSB at a starting OD_600_ of 0.1. The culture was grown at 37°C with shaking until an OD_600_ of 0.6, corresponding to approximately 1.75 × 10^9^ CFU/mL, was reached. The culture was then washed with saline and resuspended in saline to a final concentration of 1 × 10^8^ CFU/mL. Inocula were serially diluted and plated both before and after inoculation, and counts were performed to verify CFU/mL.

### Purified peptides

Human catestatin (sequence: SSMKLSFRARAYGFRGPGPQL) and mouse catestatin (sequence: RSMKLSFRTRAYGFRDPGPQL) were custom synthesized by NovoPro Bioscience at ≥95% purity. Peptides were dissolved in molecular-grade water (ThermoFisher) and stored at -80°C before thawing on ice and diluting in appropriate assay buffer.

### Wound infection model

Mice between 8-10 weeks were used. All procedures were performed under isoflurane anesthesia. Dorsal skin was shaved and depilated using hair removal cream (Veet) 24 h prior to wounding. The area was disinfected using a combination of soap, 70% ethanol and betadine. A full-thickness 6mm excisional wound was created by tracing with a 6mm biopsy punch (Integra), followed by excision with surgical scissors. Wounds were immediately photographed and treated with either 50 μM catestatin in 10 μL saline or saline alone. One h after treatment, wounds were inoculated with 10 μL of saline containing 1 × 10^6^ CFUs of MRSA strain USA300 LAC. Wounds were imaged 24 h post-infection, then excised using a 12mm biopsy punch (Integra). Each excised wound was bisected, and one half was snap-frozen in liquid nitrogen and stored at - 80°C. The other half was placed on ice before weighing, homogenizing in 1 mL PBS using a Fisherbrand 150 handheld homogenizer with a 7 mm, sawtooth-bottom stainless steel probe (Fisher Scientific), serially diluting in PBS and plating on Mannitol Salt Agar (ThermoFisher) to determine CFU/mg of tissue. Wound area was quantified using ImageJ (NIH) and expressed as a percentage of the initial wound area.

### Evans blue assay

Mice up to 4 months of age were anesthetized with isoflurane. After induction of anesthesia, mice were injected intravenously (i.v.) with 50 μL of 12.5 mg/ml Evans blue dye (Millipore Sigma) in saline. Five min later, 5 μL of 50 μM catestatin was administered by intraplantar injection in one hind paw, while 5 μL of saline was administered into the contralateral paw as a control. Paw thickness was measured by calipers (Amazon) immediately after injection and again 15 min later. Mice were then euthanized by cervical dislocation. Paw tissue was collected, dried at 50°C for 24 h, and weighed. To extract Evans blue, paws were finely chopped in 500 μL of formamide (ThermoFisher) and incubated at 50°C for 48 h. Samples were centrifuged for 15 min at 16000g, and 200 μL of supernatant was transferred to a 96-well plate. Absorbance was read at 620 nm using a Genesys 30 spectrophotometer (ThermoFisher).

### ELISA for skin samples

Skin samples were collected and snap-frozen as described above. To extract protein, samples were thawed on ice, weighed and homogenized in 1 mL of PBS containing 1% Triton X-100 (Millipore Sigma). Samples were cut into smaller pieces with surgical scissors before homogenization using a Fisherbrand 150 handheld homogenizer with 7 mm, sawtooth-bottom stainless steel probe. Samples were then freeze-thawed at -80°C and centrifuged at 10,000g for 5 min to remove cellular debris before collecting supernatants and storing at -80°C. TNF-α, CCL2 or IL-6 production was determined using the following commercial enzyme immunoassay (EIA) kits (ThermoFisher): mouse TNF alpha uncoated ELISA kit, mouse CCL2 uncoated ELISA kit, and mouse IL-6 uncoated ELISA kit. The minimum detection limits are 15.6□pg/mL for TNF-α, 15□pg/mL for CCL2, and 4 pg/mL for IL-6.

### RNA isolation from mouse skin and RT-qPCR

Mice were anesthetized using isoflurane, and dorsal hair was shaved using electric clippers and depilated. The dorsal skin was cleaned with 70% ethanol and RNaseZap™ (Fisher Scientific) prior to creating a full-thickness wound using a 6mm biopsy punch and surgical scissors as described above. The excised skin was immediately snap-frozen in liquid nitrogen and stored at - 80^°^C for use as unwounded control tissue. Three h post-injury for sterile wounds, or twenty-four h post-infection for MRSA-infected wounds, mice were euthanized by cervical dislocation, and the area around the wound was cleaned with 70% ethanol and RNaseZap™. The wound was excised using an 8mm biopsy punch (Integra) and surgical scissors, and snap-frozen in liquid nitrogen. All samples were stored at -80°C until RNA extraction. For RNA extraction, tissue was thawed on ice and 500 μL of Trizol (Millipore Sigma) was added to each sample. Tissue was cut into smaller pieces with surgical scissors, then homogenized using a Fisherbrand 150 handheld homogenizer with a 7 mm, sawtooth-bottom stainless steel probe. Next, 100 μL chloroform (Fisher Scientific) was added, tubes were shaken vigorously by hand, and then centrifuged for 15 min at 16000 g. The clear aqueous layer containing RNA was carefully collected, and RNA was purified using the Direct-zol RNA Miniprep Kit (Zymo Research) according to the manufacturer’s instructions. cDNA was generated from 1 μg f total cellular mRNA using the iScript cDNA Synthesis Kit (Bio-Rad) following the manufacturer’s instructions. qPCR was performed using 100 ng of cDNA with TaqMan Gene Expression Assays (ThermoFisher), on a QuantStudio 3 Real-Time PCR System (ThermoFisher), following the manufacturer’s instructions. The following RT-qPCR assays from ThermoFisher were used: mouse *Chga* (Mm00514341_m1), mouse *Camp* (Mm00438285_m1), mouse *Defb14* (Mm00806979_m1), mouse *S100a7a* (Mm01218201_m1), mouse *Gapdh* (Mm99999915_g1). All reactions were performed in triplicate for 40 cycles. All data are normalized to GAPDH internal controls, and no-template controls were included in all experiments. Data are expressed as fold change relative to untreated controls.

### Flow cytometry

2 cm × 1 cm sections of skin containing two 6mm MRSA-infected wounds were collected from animals and mechanically digested using surgical scissors, followed by enzymatic digestion in 10 mL RPMI (Wisent) containing 120 μg/mL Liberase TL (Millipore Sigma) and 0.01% DNase I (Millipore Sigma) with rocking for 2 h at 37°C. Digested tissue was passed through a 40μm cell strainer and spun at 300g for 7 min at 4°C. Pellets were resuspended in 3 mL ACK lysis buffer (Gibco) for 3 min to lyse the red blood cells, then diluted with 12 mL RPMI supplemented with 10% fetal bovine serum (ThermoFisher) to stop the reaction. Cells were spun at 300g for 5 min at 4°C and resuspended in 500 μL FACS buffer [1X PBS + 1% bovine serum albumin (Millipore Sigma). Cells were spun again at 300g for 5 min at 4°C, resuspended in 100 μL Fc block (Millipore Sigma) and incubated for 20 min at 4°C. Cells were spun at 300g for 5 min at 4°C and washed once with 100 μL ice-cold PBS before incubating for 15 min at room temperature in Zombie NIR™ (ThermoFisher). Cells were spun at 300g for 5 min at 4°C and washed once with 100 μL FACS buffer and then stained for 1 h at 4°C in the dark with the indicated antibodies in 100 μL FACS buffer. After staining, cells were washed twice with 100 μL FACS buffer, resuspended in 200 μL FACS buffer, and acquired on a Sony FACS SH800Z. Data were analyzed using FloReada.io analysis software. The following antibodies from Biolegend, all diluted 1:1000 in FACS buffer, were used for flow cytometry: rat anti-mouse CD45 (clone 30-F11) FITC, rat anti-mouse CD11-b (M1/70) PE/Dazzle594, rat anti-mouse Ly6C (HK1.4) APC, and rat anti-mouse (1A8) BV421. Cells were first gated to exclude debris (FSC-A vs SSC-A), then singlets were selected (SSC-A vs SSC-H). Dead cells were excluded based on their positivity for Zombie NIR. CD45^+^ cells were gated based on this population. For neutrophils and monocytes, live single cells were further gated for CD45^+^CD11b^+^ cells. Monocytes were identified as Ly6C^+^Ly6G^-^ and neutrophils as Ly6C^+^Ly6G^+^.

### Cell lines

Laboratory of Allergic Disease (LAD2) mast cells, originally derived from a male patient with mast cell leukemia^87^, were generously provided by the Metcalf lab (National Institutes of Health). Cells were cultured in StemPro-34 medium (ThermoFisher) supplemented with 2 mM L-glutamine (Wisent), 100 U/mL penicillin (Wisent), 50 μg/mL streptomycin (Wisent) and 100 ng/mL recombinant human stem cell factor (Peprotech). Cells were maintained at 37°C in 5% CO_2_ in a humidified incubator at a cell density of 0.2 × 10^6^ cells/mL. Cells were split weekly by hemi-depletion and routinely authenticated by functional responsiveness to FcεεRI-mediated activation. Human embryonic kidney (HEK)293 cells were maintained in DMEM (ThermoFisher) supplemented with 10% fetal bovine serum, penicillin-streptomycin and L-glutamine at 37°C and 5% CO_2_.

### Mast cell degranulation assay

In experiments involving biotin-conjugated IgE (bIgE), cells were first incubated overnight at 37°C and 5% CO_2_ with 0.5 μg/mL bIgE (Abbiotec). LAD2 cells were washed with HEPES buffer containing 0.4% bovine serum albumin, added to a 96-well plate at 0.025 × 10^6^ cells/well and stimulated with either HEPES+BSA buffer, 1.25, 2.5, 5 or 10 μM human catestatin, 0.5 μg/mL compound 48/80 (Millipore Sigma), or 0.025 or 0.05 μg/mL streptavidin (ThermoFisher), followed by a 30-min incubation at 37°C and 5% CO_2_. In experiments using signaling inhibitors, cells were pre-incubated with 10 μM U-73122 (Millipore Sigma), 10 μM Ro-31-8220 (Millipore Sigma), or 10 μM Wortmannin (Millipore Sigma) for 30 min at 37°C and 5% CO_2_, prior to stimulation with 10 μM catestatin or 0.5 μg/mL compound 48/80, as described above. β-hexosaminidase content in both the supernatants and cell pellets was quantified via hydrolysis of p-nitrophenyl *N*-acetyl-β-D-glucosamide (Millipore Sigma) in 0.1 M sodium citrate buffer (pH 4.5) at 37°C and 5% CO_2_ for 90 min. The reaction was terminated with 0.4 M glycine buffer (pH 10.7), and absorbance was measured at 405 nm and 595 nm using a Cytation 5 plate reader (Agilent BioTek). The release of β-hexosaminidase was calculated as a percentage of total content.

### Histamine assay

LAD2 cells were washed with BSA-free HEPES buffer, added to a 96-well plate at 0.1 × 10^6^ cells/well, and stimulated with either HEPES buffer, 1.25, 2.5, 5 or 10 μM catestatin, or 0.5 μg/mL compound 48/80. Cells were incubated at 37°C and 5% CO_2_ for 30 min. A 100 μg/mL stock solution of histamine (Millipore Sigma) was prepared and stored at -20°C. On the day of the experiment, histamine standards ranging from 500 ng/mL to 7.8 ng/mL were prepared by two-fold serial dilution. O-phthalaldehyde (OPT; Millipore Sigma) was dissolved in acetone-free methanol at 10 mg/mL and stored at 4°C in the dark until use. After incubation, supernatants and histamine standards were added to a black, flat-bottomed 96-well plate. Then, 12 μL of 1M NaOH and 2 μL of OPT were added to each well and incubated at room temperature in the dark for 4 min. Finally, 6 μL of 3M HCL was then added to stop the reaction, and fluorescence was measured with a 360 nm excitation filter and a 450nm emission filter on a Cytation 5 plate reader.

### Calcium imaging

LAD2 cells (1 × 10^6^) were allowed to adhere to sterile glass coverslips coated with 50 μg/mL fibronectin for 2 h at 37°C and 5% CO_2_. Cells were then loaded with 1:1000 Fura-2 acetoxymethyl ester (Molecular Probes) and 1:1000 Pluronic acid in HEPES buffer for 30 min in the dark at 37°C and 5% CO_2_ ^88, 89^. After two washes with HEPES buffer, cells were imaged using dual-wavelength excitation at 340 nm and 380 nm to monitor intracellular free Ca^2+^ levels. Cells were stimulated with 10 μM catestatin at the 70-second time point and with 0.5 μg/mL compound 48/80 at the 256-second time point. Calcium responses are plotted as the 340/380 fluorescence ratio over time.

### EC_50_ determination

HEK293 cells stably expressing Gα15 subunit were transfected with plasmids encoding either human MRGPRX2 or mouse Mrgprb2 cDNA. Mock-transfected cells were used as controls. Cells were seeded into 96-well plates and grown to 80-90% confluency. For Ca^2+^ mobilization assays, cells were incubated for 1 h at 37°C with the FLIPR Calcium 5 dye (Molecular Devices) in Hank’s Balanced Salt Solution (HBSS; Millipore Sigma) supplemented with 20 mM HEPES (ThermoFisher). Plates were then equilibrated at room temperature for 15 min before imaging on a FlexStation 3 (Molecular Devices). Baseline was recorded for 20 sec, peptides were added, and intracellular Ca^2+^ response was recorded for 150 sec. Data from 3-4 duplicate experiments were averaged, and EC_50_ values were calculated by normalizing to the peak response using Prism v10.0d (GraphPad).

### Molecular docking

Modelling of MRGPRX2 was achieved using cryo-EM data of MRGPRX2 bound to cortistatin 14 obtained by Yang et al (PDB:7VV4)^58^. Modelling of catestatin was achieved using nuclear magnetic resonance data of catestatin (PDB:1LV4)^59^. Docking simulations between these two structures was performed using HADDOCK2.4 web server^60, 61^, using parameters which specified the interacting residues as those defined by Yang et al. to comprise the MRGPRX2 binding pocket and the ligand binding motif^58^. The generated protein-protein complex was visualized and analyzed using the PyMOL Molecular Graphics System (Version 3.0, Schrödinger, LLC.).

### RT-qPCR for LAD2 cells

LAD2 cells (1 × 10^6^) were stimulated for 3 h with either 10 μM catestatin or 0.5 μg/mL compound 48/80 at 37°C and 5% CO_2_. Cells were centrifuged for 5 min at 200 g, the supernatant was aspirated, and the pellet was resuspended in 300 μL Trizol and stored at -20°C. RNA was purified using the Direct-zol RNA Miniprep kit, following the manufacturer’s instructions. cDNA was generated from 1 μg of total cellular mRNA using the iScript cDNA Synthesis Kit, following the manufacturer’s instructions. qPCR was performed using 100 ng of cDNA with TaqMan Gene Expression Assays (ThermoFisher) on a QuantStudio 3, following the manufacturer’s instructions. The following RT-qPCR assays from ThermoFisher were used: human *Tnf* (Hs00174128_m1), human *Ccl2* (Hs00234140_m1), human *Il6* (Hs00174131_m1), human *Il3* (Hs00174117_m1), human *Csf2* (Hs00929873_m1) human *Gapdh* (Hs02786624_g1). All reactions were performed in triplicate for 40 cycles. All data are normalized to GAPDH internal controls, and no-template controls were included in each experiment. Data are expressed as fold change relative to untreated controls.

### ELISA for LAD2 cells

LAD2 cells (1 × 10^6^) were stimulated for either 3 h or 24 h with either 10 μM catestatin or compound 48/80 (0.5 μg/mL or 1μg/mL) at 37°C and 5% CO_2_. Cell-free supernatants were collected and analyzed for prostaglandin D_2_ (PGD_2_), TNF-α or CCL2 production using the following commercial enzyme immunoassay (EIA) kits: Prostaglandin D_2_ EIA Kit (Cayman Chemicals), Human TNF-alpha uncoated ELISA Kit (ThermoFisher), Human CCL2 uncoated ELISA Kit (ThermoFisher) and Human GM-CSF uncoated ELISA Kit (ThermoFisher). The minimum detection limits are 19.5 pg/mL for PGD_2_, 4 pg/mL for TNF-α, 7 pg/mL for CCL2 and 6 pg/ml for GM-CSF.

### Western blots

WT and MRGPRX2-knockout (X2-KO) LAD2 cells (1 × 10^6^) were washed and resuspended in HEPES buffer before stimulation with either 10 μM catestatin or 0.5 μg/mL compound 48/80 at 37°C and 5% CO_2_ for the indicated timepoints. Cells were then centrifuged at 2500g for 5 min at 4°C, and pellets were resuspended in RIPA buffer (Millipore Sigma) containing a Pierce protease and phosphatase inhibitor cocktail (ThermoFisher). Lysates were rocked for 15 min, then centrifuged at 14000g for 15 min at 4°C. Total protein concentrations in the supernatants were quantified using the Pierce BCA assay kit (ThermoFisher), as per the manufacturer’s instructions, and samples were stored at -80°C. A total of 100 μg of protein was separated on 6-12% polyacrylamide gradient gels and transferred onto Amersham Protran 0.45μm nitrocellulose membranes using a semi-dry transfer system. Membranes were blocked in 4% BSA in TBS-T (1X Tris-buffered saline supplemented with 0.1% Tween-20) for 1 h at room temperature, followed by three washes with TBS-T. Membranes were then incubated overnight at 4°C with 1:1000 primary P-p44/42 MAPK (T202/Y204) Rabbit mAb (Cell Signaling Technology, Cat. No. 4370S). After washing (twice with TBS-T, once with TBS), membranes were then incubated with 1:2000 HRP-conjugated anti-rabbit secondary antibody (Cell Signaling Technology, Cat. No. 7074P2). for 1 h at room temperature. Bands were visualized using a 5-min incubation with Clarity Western ECL Substrate (Bio-Rad) and imaged with ChemiDoc system (Bio-Rad). After imaging, the membranes were stripped with stripping buffer for 30 min at room temperature, re-blocked in 1X 4% BSA in TBS-T for 1 h at room temperature, and washed three times with TBS-T. Membranes were subsequently incubated overnight at 4°C with 1:1000 primary p44/42 MAPk (Erk1/2) Mouse mAb (Cell Signaling Technology, Cat. No. 9107S), followed by 1-h incubation with 1:2000 HRP conjugated anti-mouse secondary antibody (Cell Signaling Technology, Cat. No. 7076P2). at room temperature. After washing three times, bands were again developed using Clarity Western ECL Substrate and imaged using the ChemiDoc.

### Lactate dehydrogenase assay

LAD2 cells (5 × 10^3^) were stimulated with 10 μM catestatin for 30 min at 37°C and 5% CO_2_. Following treatment, manufacturer’s instructions were followed for the CyQUANT LDH Cytotoxicity Assay (ThermoFisher).

### Statistics

Data are presented as mean ± standard error of mean (SEM) and were analyzed by two-tailed unpaired Student’s *t*-test or two-way ANOVA, as specified, using Prism v10.0d (GraphPad). Statistical significance was defined as *P* < 0.05.

## ACKNOWLEDGEMENTS

The authors thank Dr. Nathachit Limjunyawong (Mahidol University) for help with EC_50_ analyses and reagents. USA300 LAC was kindly provided by Dr. Georgina Cox (University of Guelph). LAD2 cells were generously provided by the Metcalf lab at NIAID. Mrgprb2-KO mice were generated in Dr. Xinzhong Dong’s lab at Johns Hopkins University. We also thank Dr. Xintong Dong University of Texas at Dallas) for critically reviewing the manuscript.

## FUNDING

This work was supported by the J. P. Bickell Foundation (Grant #056063), the Canadian Foundation for Innovation – John R. Evans Leader Fund for Innovation (Grant #42143), and the College of Biological Science at the University of Guelph (Grant #055519).

## CREDIT AUTHORSHIP CONTRIBUTION STATEMENT

**Colin Guth:** Writing – review & editing, Writing – original draft, Formal analysis, Data curation, Conceptualization. **Hannah Dychtenberg:** Formal analysis, Data curation. **Erin Rudolph:** Data curation. **Austin Pozniak:** Data curation**. Sukhmeen Gill:** Data curation. **Priyanka Pundir:** Writing – review & editing, Writing – original draft, Supervision, Project Administration, Funding Acquisition, Formal analysis, Data curation, Conceptualization.

## DECLARATION OF COMPETING INTEREST

The authors declare that they have no known competing financial interests or personal relationships that could have appeared to influence the work reported in this paper.

## SUPPLEMENTARY INFORMATION

Supplementary data are available for this article.

## DATA AVAILABILITY

The original contributions presented in this study are included in the article/supplementary material. Further inquiries can be directed to the corresponding author.

## ABBREVIATIONS

B2-/-: Mrgprb2-knockout
bIgE: Biotin-conjugated IgE
BSA: Bovine serum albumin
CCL2: C-C motif ligand 2
CFU: Colony forming unit
Cryo-EM: Cryo-electron microscopy
CST: Catestatin
Defb14: Mouse beta-defensin 14
GM-CSF: Granulocyte-macrophage colony-stimulating factor
GPCR G: protein-coupled receptor
HEK293: Human embryonic kidney
IL: Interleukin
KO: Knockout
LAD2: Laboratory of Allergic Diseases 2
LDH: Lactate dehydrogenase
Mrgprb2: Mas-related G protein-coupled receptor b2
MRGPRX2: Mas-related G protein-coupled receptor X2
MRSA: Methicillin-resistant *Staphylococcus aureus*
PGD_2_: Prostaglandin D_2_
PI3K: Phosphatidylinositol 3-kinase
PKC: Protein kinase C
PLC: Phospholipase C
TNF: Tumor necrosis factor-α
WT: Wildtype
X2-KO: MRGPRX2-knockout

## Notes

### Competing Interest Statement

The authors have declared no competing interest.

