## Supplementary Data for "The neuroendocrine peptide catestatin promotes clearance of cutaneous *Staphylococcus aureus* through mast cell Mrgpr activation"

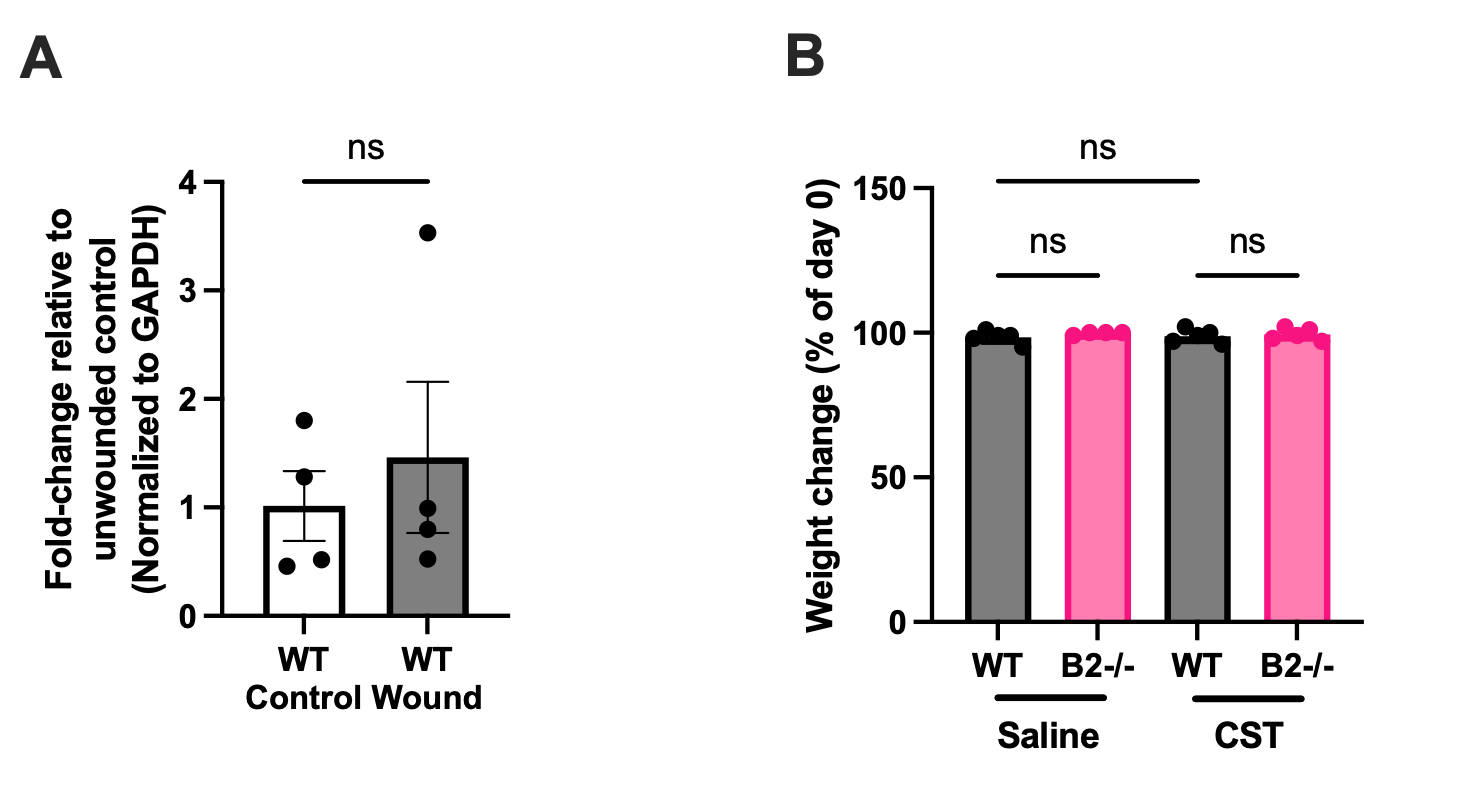

**Figure S1. Full-thickness cutaneous wounds do not upregulate endogenous catestatin.**
**(A)** mRNA expression of chromogranin A in the skin of wildtype (WT) C57Bl/6 mice, 3 h post-wounding, normalized to GAPDH and expressed relative to uninjured skin. n=4.
**(B)** Change in body weight 24 h post-infection in saline- or catestatin (CST)-treated WT (n = 5 each), and saline- or CST-treated Mrgprb2-KO (B2-/-) mice (n = 5 and 4, respectively), expressed as percentage relative to pre-wound weight.
Data are presented as mean ± SEM and analyzed by unpaired Student’s t-test or one-way ANOVA.

Related to Figure 1

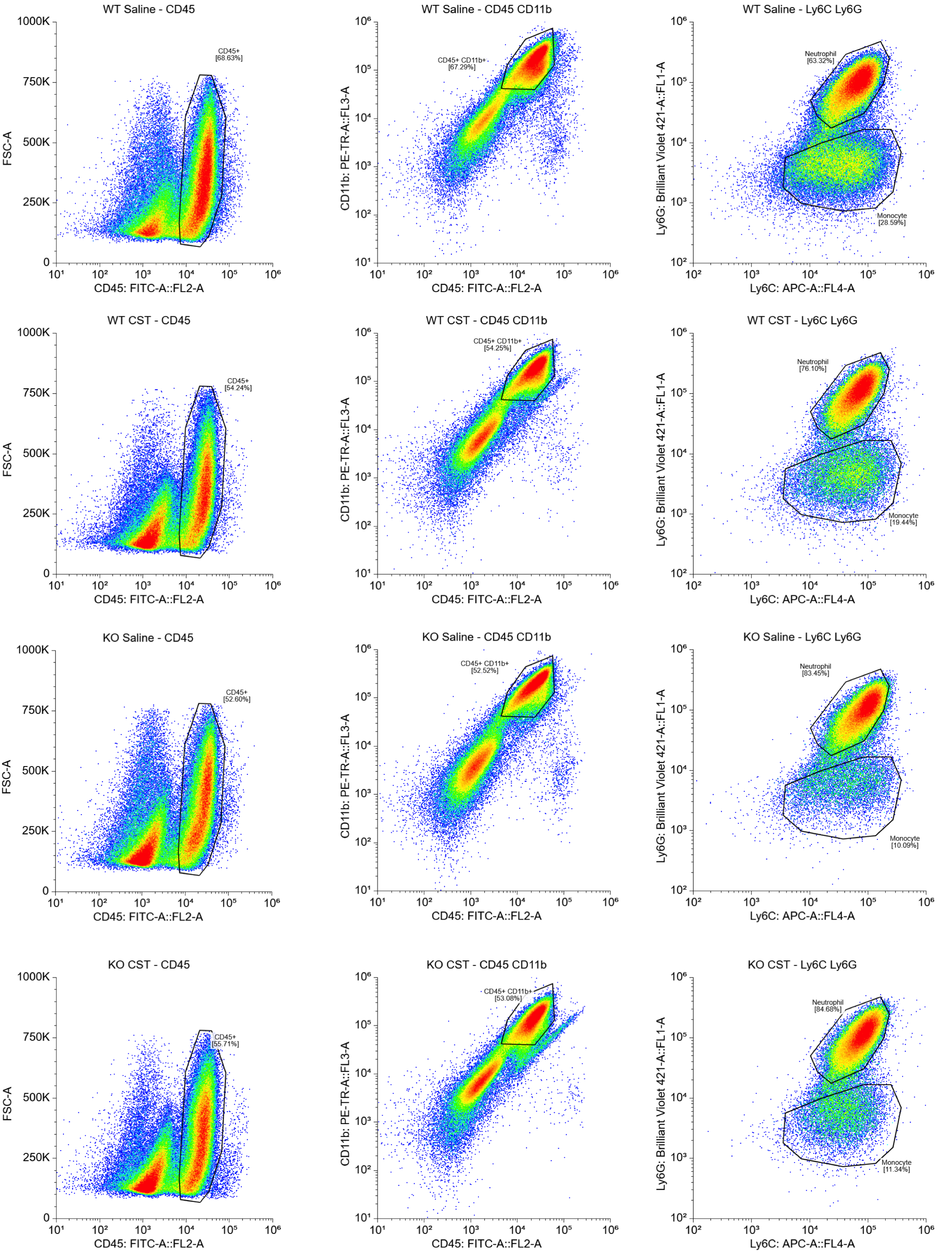

**Figure S2. Gating strategy for flow analysis of skin samples.**

Representative flow cytometry plots illustrating the sequential gating strategy used to identify leukocyte populations in wound tissue following MRSA infection. Debris was excluded using FSC-A vs SSC-A, and singlets were gated via SSC-A vs SSC-H. Zombie NIR was used to exclude dead cells. Live CD45⁺ leukocytes were identified, and CD11b⁺ cells were further analyzed to distinguish neutrophils (Ly6C⁺Ly6G⁺) and monocytes (Ly6C⁺Ly6G⁻).

Related to Figure 2

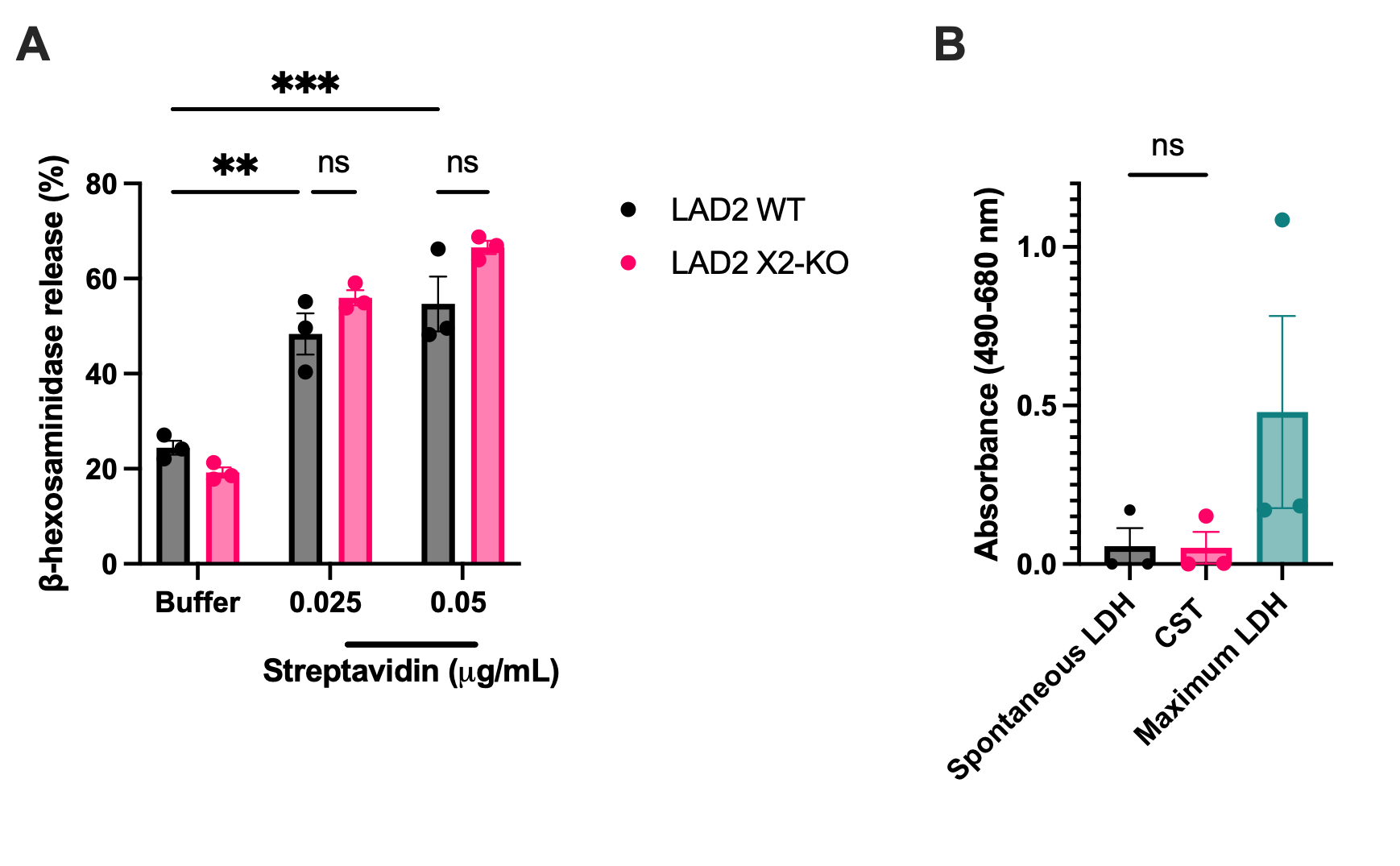
**Figure S3. Degranulation and cytotoxicity in LAD2 cells**.

**(A)** β-hexosaminidase release from biotin-conjugated IgE (bIgE)-sensitized WT and MRGPRX2-knockout (X2-KO) LAD2 cells stimulated with 0.025 and 0.05 µg/ml streptavidin. n=3.

**(B)** LDH release from LAD2 cells in response to 10 µM catestatin (CST). n=3.

Data are expressed as mean ± SEM, analyzed by unpaired Student’s *t*-test or one-way ANOVA. **P<0.01, ***P<0.001.

Related to Figure 4

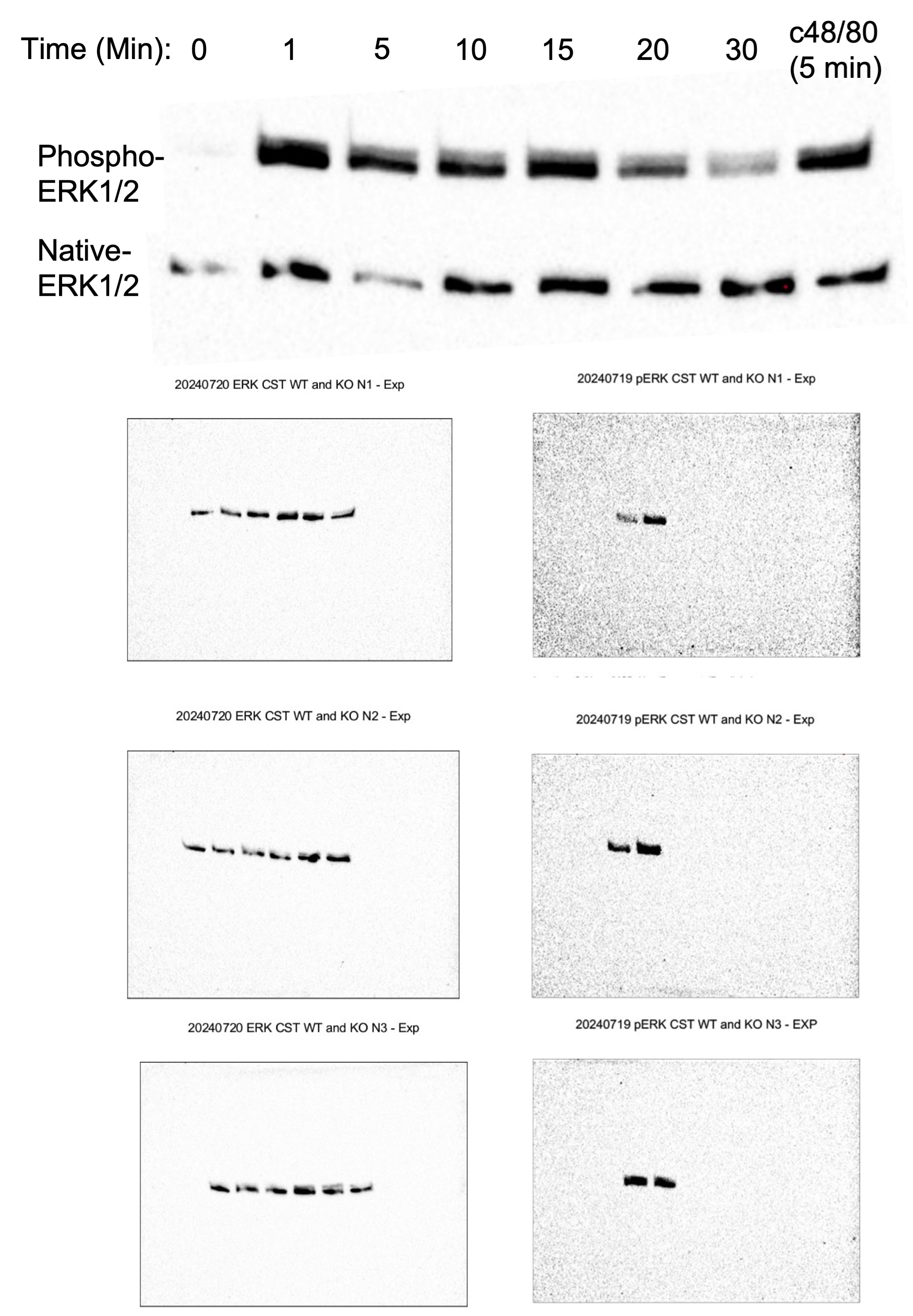
**Figure S4. Time course of ERK1/2 phosphorylation induced by catestatin.**

Western blot images of native and phosphorylated ERK1/2 from LAD2 cells stimulated with catestatin (10 µM) for different durations. Compound 48/80 (10 µg/mL) is included as a positive control.

Related to Figure 6
